# A Comparison of Temporal Response Function Estimation Methods for Auditory Attention Decoding

**DOI:** 10.1101/281345

**Authors:** Daniel D.E. Wong, Søren A. Fuglsang, Jens Hjortkjær, Enea Ceolini, Malcolm Slaney, Alain de Cheveigné

## Abstract

The decoding of selective auditory attention from noninvasive electroencephalogram (EEG) data is of interest in brain computer interface and auditory perception research. The current state-of-the-art approaches for decoding the attentional selection of listeners are based on temporal response functions (TRFs). In the current context, a TRF is a function that facilitates a mapping between features of sound streams and EEG responses. It has been shown that when the envelope of attended speech and EEG responses are used to derive TRF mapping functions, the TRF model predictions can be used to discriminate between attended and unattended talkers. However, the predictive performance of the TRF models is dependent on how the TRF model parameters are estimated. There exist a number of TRF estimation methods that have been published, along with a variety of datasets. It is currently unclear if any of these methods perform better than others, as they have not yet been compared side by side on a single standardized dataset in a controlled fashion. Here, we present a comparative study of the ability of different TRF estimation methods to classify attended speakers from multi-channel EEG data. The performance of the TRF estimation methods is evaluated using different performance metrics on a set of labeled EEG data from 18 subjects listening to mixtures of two speech streams.

## 1 INTRODUCTION

A fundamental goal of auditory neuroscience is to understand the mapping between auditory stimuli and the cortical responses they elicit. In magneto/electro-encephalography (M/EEG) studies, this mapping has predominantly been measured by examining the average cortical evoked response potential (ERP) to a succession of repeated short stimuli. More recently, these methods have been extended to continuous stimuli such as speech by using linear stimulus-reponse models, broadly termed ‘temporal response functions’ (TRFs). The TRF characterizes how a unit impulse in an input feature corresponds to a change in the M/EEG data. TRFs can be used to generate continuous predictions about M/EEG responses or stimulus features, as opposed to characterizing the response (ERP) to repetitions of the same stimuli. Importantly, it has been demonstrated that the stimulus-response models can be extracted both from EEG responses to artificial sound stimuli (16) but also from EEG responses to naturalistic speech (17). A number of studies have considered mappings between the slowly varying temporal envelope of a speech sound signal (*<*10 Hz) and the corresponding filtered M/EEG response (16, 28, 11, 12). However, TRFs are not just limited to the broadband envelope, but can also be obtained with the speech spectrogram (9, 10), phonemes (8), or semantic features (4). This has opened new avenues of research into cortical responses to speech, advancing the field beyond examining responses to repeated isolated segments of speech.

TRF decoding methods have proven particularly apt for studying how the cortical processing of speech features are modulated by selective auditory attention. A number of studies have considered multitalker ‘cocktail party’ scenarios, where a listener attends to one speech source and ignores others. It has been demonstrated that both attended and unattended acoustic features can be linearly mapped to the cortical response (9, 10, 28, 29, 38), or, conversely, from the cortical response to the speech features (23, 20, 14, 9, 10, 19. Differences in the accuracy of TRF-derived predictions between the attended and unattended speech signal can be used to predict or ‘decode’ to whom a listener is attending based on unaveraged M/EEG data. Single-trial measures of auditory selective attention in turn suggests BCI perspectives, for instance, for hearing instrument control.

The ability of TRF models to generalize to new data is generally limited by the need to estimate a relatively large number of parameters based on noisy single-trial M/EEG responses. Like many aspects of machine learning, this necessitates regularization techniques that constrain the TRF model coefficients to prevent overfitting. A number of methods for regularizing the TRF have been presented in various studies. Each of these methods attempt to address the challenge of having sufficient data to compute a reliable TRF function. To reduce the data requirement, regularization can be applied in the form of a smoothness and/or sparsity constraint.

To date, little work has been done to compare these methods against each other. A meta-analysis would be difficult as many variables, such as subjects, stimuli and data processing are different between each study. The present paper proposes a standardized dataset, based on the attended-versus-unattended talker discrimination task, as well as preprocessing and evaluation procedures to compare these algorithms. In addition, the present paper examines the relationship between different evaluation metrics to highlight their similarities and differences. The TRF methods have been implemented in the publicly available Telluride Decoding Toolbox^1^.

## 2 MATERIAL AND METHODS

Temporal response functions can be used to predict the EEG response to a multi-talker stimulus from the attended speech envelope or, alternatively, to reconstruct the attended speech envelope from the EEG response. The first case is denoted as a “forward TRF” (as it maps from speech features to neural data) and the second as a “backward TRF” (as it maps from neural data back to speech features).

### 2.1 Temporal Response Functions

The TRF methods described below map a matrix **X** = [*x*_(*t*, *f*), *c*_] to a matrix **Y** = [*y*_*t*_]:

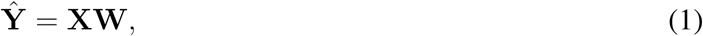

where 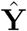 is the TRF model prediction in the form of a time-dimension *t* vector, and **X** is the TRF model input matrix with time-dimension *t* and channel-dimension *c*. **X** is augmented to include time-lagged versions of the data with a limited range of time lags, for example −500 ms to +500 ms, so that the model can handle delays and convolutional mismatch between **X** and **Y**. These time lags are denoted as dimension *f* and are combined with the time dimension *t* to form a single dimension when performing matrix multiplications. For a forward TRF model, **X** is a representation of the stimulus (e.g. single-channel speech envelope) and **Y** is the EEG response. In this case, a TRF can be computed for each EEG electrode channel. For a backward TRF model, **X** is the EEG data with channel dimension *c* and **Y** is a representation of the stimulus.

In the following subsections we introduce different approaches to estimating the linear TRF model parameters, **W**. Each method uses different regularization techniques to optimize the generalizability of the mapping functions.

#### 2.1.1 Ordinary Least Squares (OLS)

The TRF filter coefficients can be estimated via ordinary least squares:

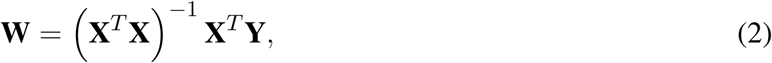

where **X**^*T*^**X** is the estimated covariance matrix and **X**^*T*^**Y** is the estimated cross-covariance matrix. The ordinary least-squares solution was here estimated using the Cholesky decomposition method, via the *mldivide* routine in Matlab. One advantage of the OLS estimator is that it has no additional hyperparameters that must be optimized. However, in practice the OLS estimator is often outperformed by the regularized solutions described in the following subsections. This is often the case when the regressor, **X**, is highdimensional, has highly correlated columns and has a poorly estimated covariance matrix given limited amounts of training data.

#### 2.1.2 Ridge

Ridge regression minimizes the residual sum of squares, but puts an *L*2 constraint on the regression coefficients which biases the solution. Ridge regression corresponds to imposing a Gaussian prior on the filter coefficients (37). The ridge solution is:

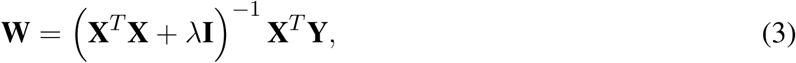

where *λ* is the regularization parameter that controls the amount of parameter shrinking.

#### 2.1.3 Low-Rank Approximation (LRA)

The LRA-based regression relies on a low-rank approximation of the covariance matrix, **X**^*T*^ **X**. This is achieved by employing a singular value decomposition (SVD) of **X**^*T*^ **X**:

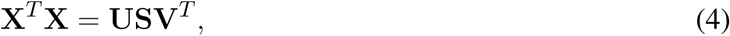

where **U** and **V** are orthonormal matrices that contain respectively the left and right singular vectors, and where **S** is a diagonal matrix, **S** = diag(*s*_1_, *s*_2_,.*.s*_*d*_) with sorted diagonal entries. Since **X**^*T*^ **X** is a positive semidefinite matrix we have **U** = **V**. LRA uses a rank-*K* approximation of **X**^*T*^ **X** by only retaining the first 1 *≤ K ≤ d* diagonal elements of **S**. By forming 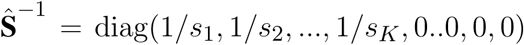, the regression coefficients can be estimated from:

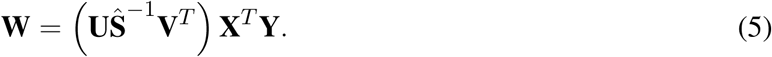

The number of diagonal elements, *K*, to retain are typically chosen such that a diagonal element is retained if the sum of the eigenvalues to be kept cover a fraction *λ* of the overall sum, or 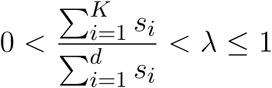 Note that the regularization parameter, *λ*, here is analogous to *λ* for Ridge Regression, but that the values are not comparable between the two.

#### 2.1.4 Shrinkage

Shrinkage (3, 13) is a method used for biasing the covariance matrix by flattening its eigenvalue spectrum with some tuning parameter, *λ*. In the context of regression, the Shrinkage solution is

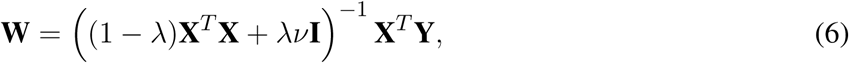

where *ν* is here defined as the average eigenvalue trace of the covariance matrix (X_*T*_X). When *λ* = 0, it becomes the standard ordinary least squares solution. When *λ* = 1, the covariance estimator becomes diagonal (i.e. it becomes spherical) (3).

These regularization schemes are related. Whereas Ridge Regression and Shrinkage both penalize extreme eigenvalues in a smooth way, LRA discards eigenvalues. Ridge and Shrinkage in other words flatten out the eigenvalue trace. Ridge shifts it up, and Shrinkage shrinks it towards an average value *ν*(3), whereas LRA cuts if off.

#### 2.1.5 Tikhonov

Tikhonov regularization takes advantage of the fact that there is usually a strong correlation between adjacent columns of **X** when **X** includes time shifts, because of the strong serial correlation of the stimulus envelope (for the forward model) or the filtered EEG (for the backward model). In other words, Tikhonov regularization imposes *temporal smoothness* on the TRF. While Ridge Regression is a special type of Tikhonov regularization, the scheme which we shall refer to as *Tikhonov regularization* achieves temporal smoothness by putting a constraint in the derivative of the filter coefficients (17, 18, 15). Here we focus on first order derivatives of the filter coefficients and assume that the first derivatives can be approximated by 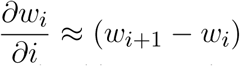 for any neighboring filter pairs *w*_*i*+1_ and *w*_*i*_. Tikhonov regularized TRF filters can, under this approximation, be implemented as:

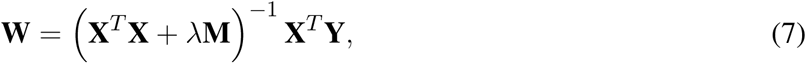

where

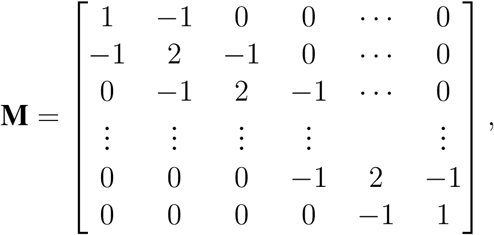

Note that cross-channel leakage can occur whenever the regressor, **X**, reflects data recorded from multiple channels. This means that filter endpoints can be affected by neighboring channels as a result of the offdiagonal elements in the **M** matrix. However, as long as the TRFs have sufficiently long memory, it is likely that the filter values at the endpoints will attain low values, such that the cross-channel leakage effects become negligible.

#### 2.1.6 Elastic Net

Whereas the aforementioned regularization techniques often show improvements over the ordinary least regression in terms of generalizability, they tend to preserve all regressors in the models. This can e.g. result in nonzero filter weights assigned to irrelevant features. Lasso regression attempts to overcome this issue by putting an L1-constraint on the regression coefficients (32). This serves to drive unnecessary coefficients in the TRF towards zero. Lasso has been found to perform well in many scenarios, although it was empirically demonstrated that it is outperformed by Ridge regression in nonsparse scenarios with highly correlated predictors (32, 39). In such scenarios, *Elastic Net* regression (39) has been found to improve the predictive power of Lasso by combining Lasso with the grouping effect of Ridge regression. The elastic net has two hyperparameters: *α* controlling the balance between L1 (lasso) and L2 (ridge) penalties, and *λ* controlling the overall penalty strength. For the purpose of this paper, we use a readily available algorithm, GLMNET (30), for efficiently computing the elastic net problem. This is a descent algorithm for solving the following problem:

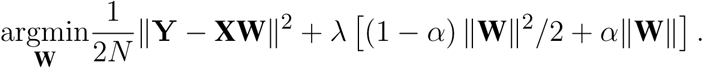

### 2.2 Evaluating Performance

#### 2.2.1 Characterizing TRF Model Fit

While the objective function of linear TRFs is minimizing the mean-squared-error, the goodness of fit is typically analyzed in terms of Pearson’s correlation between predicted and actual values due to the difference in dimensionality between EEG and audio data. The term *regression accuracy* will henceforth be used to characterize the goodness of fit for TRF models trained and evaluated on attended audio features (*r*_*attended*_). For forward TRF models, regression accuracies were measured by the Pearson’s correlation between the actual EEG and the EEG predicted by the attended envelope over the test folds. This was done separately for each EEG channel. Similarly, for backward TRF models, regression accuracies were measured by the correlation between the attended envelope and its EEG-based reconstruction. Other metrics for assessing the predictive performance of the TRF models have been previously proposed (31). However, for simplicity and to be consistent with previous studies (23, 9, 10), this paper characterizes the goodness of the fit using Pearson’s correlation coefficients.

In the forward case, multiple EEG channels are predicted by the TRF. Rather than using multiple correlation coefficients to characterize the regression accuracy in this case, we chose to take the average of the correlation coefficients between the predicted channels and the actual EEG data as a validation score. The assumption with this approach is that low correlation scores will cancel out. We used the same metric over the test set to characterize the fit of the TRF. In the backward case, characterizing the fit is straightforward as the TRF predicts a single audio envelope that can be correlated with the attended audio envelope.

#### 2.2.2 Decoding Selective Auditory Attention With TRF Models

Performance was also evaluated on a classification task based on the TRF model. The task of the classifier was to decide, on the basis of the recorded EEG and the two simultaneous speech streams presented to the listener (see Section 2.4), to which stream the subject was attending. The classifier had to make this decision on the basis of a segment of test data, the duration of which was varied as a parameter (1, 3, 5, 7, 10, 15, 20 and 30s), which will be referred to as the decoding segment length. This duration includes the kernel length of the TRF (500 ms). The position of this interval was stepped in 1s increments.

As described further in section 2.2.3, a nested cross-validation loop was used to tune the regularization parameter (where applicable) and test the trained classifier on unseen data. In the outer cross-validation loop the data were split into training/validation (90%) and test (10%) sets. In the inner cross-validation loop the regularization parameter was tuned (where applicable) and the TRF trained on the training/validation set, after which the trained TRF was tested on the test set. Using this TRF model, the classification relied on correlation coefficients between the attended audio and the EEG, and between the unattended audio and the EEG. These correlation coefficients were computed over the aforementioned restricted time window. These coefficients were used to classify whether the subject was attending to one stream or the other. For a backward TRF model, classification hinged merely on which correlation coefficient was largest (stream A or stream B). Performance of this classifier was evaluated on the test set. For a forward TRF model, the situation is more complex because there is one TRF model per EEG channel. For each of the 66 channels a pair of correlation coefficients was calculated (one each for unattended and attended streams), and this set of pairs was used to train a support vector machine (SVM) classifier with a linear kernel and a soft margin constant of 1. This training was performed on the training/validation set and the classifier was applied to the test set.

The training/testing process was repeated with the 9 other train/test partitions and the score averaged over all 10 iterations. In every case, the classifier trained over the entire training/validation set was tested on a short interval of data, the duration of which was varied as a parameter, as explained above. An illustration of this classification task is shown in figure 1.

**Figure 1.**
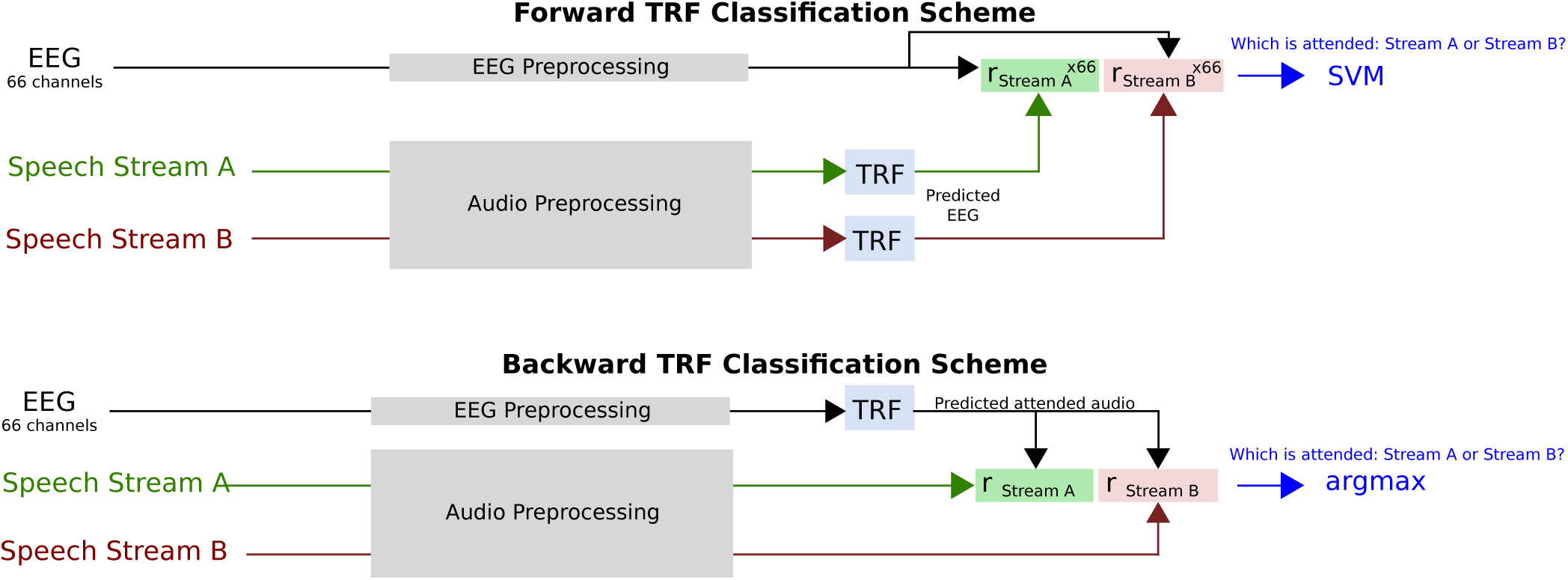
Diagram of classification task. For the forward TRF, 66 EEG channels are predicted from the speech stream A and B envelopes. After correlation with the 66 channel EEG data, this results in 66 correlation coefficients for each speech stream, which are used as features for the SVM to distinguish the attended talker. For the backward TRF, a single attended audio envelope channel is estimated from the EEG data. After correlation with the speech stream A and B envelopes, a single correlation coefficient for each speech stream is obtained. Classification of the attended talker is performed by determining the larger coefficient.

Classification performance was characterized for different decoding segment durations using the raw classification score, receiver operating characteristic (ROC) curve, and information transfer rate (ITR). The raw classification score measured what proportion of trials were classified correctly. It should be noted that in measuring classification performance, the two classes were balanced. The ROC curve characterizes the true-positive and false-positive rates for decoding segment trials where the classifier discrimination function lies above a given threshold, as the threshold is varied. The ITR metric corresponds to the number of classifications that can be reliably made by the system in a given amount of time. The dependency of ITR on decoding segment length is a tradeoff between two effects. On one hand, longer decoding segments allow more reliable decisions. On the other, short durations allow a larger number of independent decisions. There is thus an optimal decoding segment duration. A number of metrics to compute the ITR have been proposed. The most common is the Wolpaw ITR (36), which is calculated in bits per minute as:

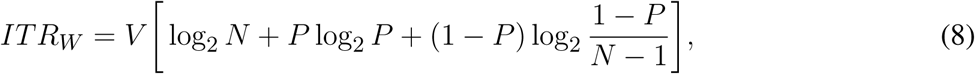

where *V* is the speed in trials per minute, *N* is the number of classes, and *P* is the classifier accuracy. We also report the Nykopp ITR, which assumes that a classification decision does not need to be made on every trial (21). This can be done by first calculating the confusion matrix *p* for classifier outputs where the classifier decision function exceeded a given threshold. This threshold is adjusted to maximize:

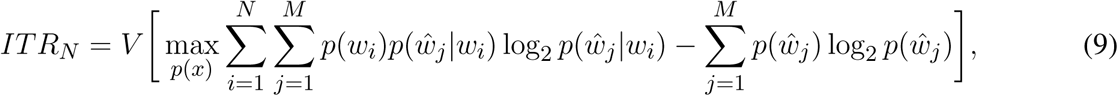

where *p*(*w*_*i*_) is the probability of the actual class being 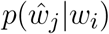 is the probability of the predicted class being class *j* given the actual class being class *i*, and 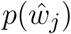 is the probability of the predicted class being class *j*.

#### 2.2.3 Cross-Validation Procedure

The TRF models were all trained and tested using cross-validation with a 10-fold testing procedure involving nested cross-validation loops. During this cross-validation procedure the TRFs were characterized under a N-fold testing framework where the data was divided into 10 folds. One fold was held out for testing, while data from the remaining 9 folds were used to compute the TRF. An additional cross-validation loop on the remaining 9 folds was used to tune the hyperparameters. In this cross-validation, the regularization parameter was adjusted to maximize the correlation coefficient between the TRF model prediction and the actual measured data. For Ridge and Lasso regularization schemes that allowed a regularization parameter between zero and infinity, a parameter sweep was performed between 10^−6^ and 10^8^ in 54 logarithmically-spaced steps. This was done using the following formula:

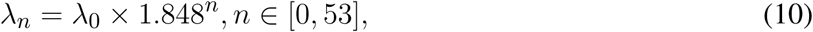

where *λ*_0_*≡*10^−6^. For LRA, Elastic Net, and Shrinkage schemes, where the regularization parameter range was between 0 and 1, a parameter sweep was performed between 10^−6^ and 1 using a log-sigmoid transfer function that compresses the values between 0 and 1 using the following iterative formula:

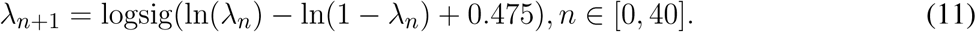

The weights of the TRF models generated for each inner cross-validation fold were then averaged to generate an overall cross-validated model that could then be applied to the test set.

### 2.3 Implementation

The implementations of the TRF algorithms used here are distributed as part of the Telluride Decoding Toolbox^2^, specifically in the FindTRF.m function of that toolbox. Data preprocessing, TRF model training, and evaluation were implemented with the COCOHA Matlab Toolbox^3^.

### 2.4 Stimuli

A previous report gives a detailed description of the stimuli and data collection procedure (14). In brief, a set of speech stimuli were recorded by one male and one female professional Danish speakers speaking different fictional stories. These recordings were performed in an anechoic chamber at the Technical University of Denmark (DTU). The recording sampling rate was 48 kHz. Each recording was divided into 50-s long segments for a total of 65 segments.

### 2.5 Experimental Procedure

The 50-s long speech segments were used to generate auditory scenes comprising a male and a female simultaneously speaking in anechoic or reverberant rooms. The two concurrent speech streams were normalized to have similar root-mean square values. The speech stimuli were delivered to the subjects via ER-2 insert earphones (Etymotic Research). The speech mixtures were presented binaurally to the listeners, with the two speech streams lateralized at respectively *-*60° and +60° along the azimuth direction and a source-receiver distance of 2.4 meters. This was achieved using nonindividualized head-related impulse responses that were simulated using the room acoustic modeling software, Odeon (version 13.02). Each subject undertook sixty trials in which they were presented the 50s-long speech mixtures. Before each trial, the subjects were cued to listen selectively to one speech stream and ignore the other. After each trial, the subjects were asked a comprehension question related to the content of the attended speech stream. The position of the target streams as well as the gender of the target speaker were randomized across trials. Moreover, the type of acoustic room condition (either anechoic, mildly reverberant or highly reverberant) were pseudo-randomized over trials. In the analysis, data recorded from all acoustic conditions were pooled together.

### 2.6 Data Collection

Electroencephalography (EEG) data were recorded from 19 subjects in an electrically shielded room while they were listening to the stimuli described above. Data from one subject were excluded from the analysis due to missing data from several trials. The data were recorded using a Biosemi Active 2 system, with a sampling rate of 512 Hz. Sixty-four channel EEG data (10/20-system) were recorded from the scalp. Six additional electrodes were used for recording the EEG at the mastoids, and vertical and horizontal electrooculogram (V and H-EOG). Approximately 1 hour of EEG data was recorded per subject. This study was carried out in accordance with the recommendations of ‘Fundamental and applied hearing research in people with and without hearing difficulties, Videnskabsetiske komitee’. The protocol was approved by the Science Ethics Committee for the Capital Region of Denmark. All subjects gave written informed consent in accordance with the Declaration of Helsinki.

### 2.7 Data Preprocessing

#### 2.7.1 EEG Data

50 Hz line noise and harmonics in the EEG data were filtered out by convolution with a 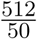 sample square window (the non-integer window size was implemented by interpolation) (5). The EEG data was then downsampled to 64 Hz using a resampling method based on the Fast Fourier Transform (FFT). A 1st order detrend was performed on the EEG data to minimize filter startup artifacts. EEG data were highpassed at 0.1 Hz using a 4th order forward-pass Butterworth filter. The group delay was less than 2 samples above 1 Hz.

The joint decorrelation framework (6) was employed to remove eye artifacts in an automated fashion. Let **X** = [*x*_*tj*_] be a matrix that contains EEG data from each electrode, *j*, for each time sample *t*. In this implementation, a conservative eye artifact time-point detection was first performed by computing a Z-score on 1-30 Hz bandpassed VEOG and HEOG bipolar channels and marking time samples where the absolute Z-score on either channel exceeded 4. This is similar to the eyeblink detection method implemented in the FieldTrip EEG processing toolbox (22). This resulted in a subset of time samples, *A*, indexing the temporal locations of each EOG artifact. An artifact covariance matrix **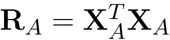** was then computed from the EEG (and EOG) data, **X**_*A*_ = [*x*_*aj*_], at the artifact time samples *a ∈ A*. The generalized eigenvalue problem was then solved for **R**_*A*_**v** = *λ***Rv**, where **R** = **X**^*T*^ **X** is the covariance matrix for the entire EEG dataset. The resulting eigenvectors **V**, sorted by eigenvalue, explain the maximum difference in variance between the artifact and data covariance matrices. Components corresponding to eigenvalues *>* 80% of the maximum eigenvalue were regressed out of the data. In practice, this 80% threshold is a conservative one, typically resulting in the removal of one or two components. Lastly, the EOG channels were removed from the data, which was then referenced to a common average over all channels.

For the TRF analysis, the EEG was bandpassed between 1-9 Hz using a windowed sync type I linear-phase finite-impulse response (FIR) filter, shifted by its group delay to produce a zero-phase (35) with a conservatively chosen order of 128 in order to minimize ringing effects. This frequency range was selected as it has been shown that cortical responses time-lock to speech envelopes in this range (23). As part of the cross-validation procedure, individual EEG channels were finally centered and standardized (Z-normalized) across the time dimension using the mean and standard deviation of the training data. A kernel length of 0.5 s (33 samples) was used when computing the TRFs.

#### 2.7.2 Audio Features

The TRF estimation methods used for attention decoding attempt to characterize a relationship between features of attended speech streams and EEG activity. We calculated temporal envelope representations from each of the clean speech streams (i.e. without reverberation). We did not try to derive them from the reverberant or mixed audio data, as explored elsewhere (14, 1). In trials with reverberant speech mixtures, we used envelope representations of the underlying clean signals to estimate the TRFs. To derive the envelope representations, we passed monaural versions of both attended and unattended speech streams through a gammatone filterbank (26). The envelope of each filterbank output was calculated via the analytic signal obtained with the Hilbert transform, raised to the power of 0.3. This rectification and compression step was intended to partially mimic that which is seen in the human auditory system (27). The audio envelope was then calculated by summing the rectified and compressed filterbank outputs across channels. The audio envelope data was subsequently downsampled to the same sampling frequency as the EEG (64 Hz) using an FFT-based resampling method. The EEG and envelopes were then temporally aligned using start-trigger events recorded in the EEG. The envelopes were subsequently lowpassed at 9 Hz. As part of the cross-validation procedure, audio envelopes were finally centered and standardized (Z-normalized) across the time dimension using the mean and standard deviation of the attended speech envelope in the training data.

### 2.8 Statistical Analysis

All statistical analyses were calculated using MATLAB. Repeated-measures analysis of variance (ANOVA) tests were used to assess differences between the regression accuracies (section 2.2.1) and classification performances 2.2.2 obtained with the different TRF estimation methods. Regression accuracies and classification performances for individual subjects were averaged across folds prior to statistical comparison.

Given the non-Gaussian distribution of regression accuracies (range −1 to 1) and classification performance metrics (range 0 to 1), Fisher Z-transforms and arcsine transforms were applied to these measures, respectively, prior to statistical tests and correlations.

## 3 RESULTS

The TRF estimation methods introduced in Section 2 were used to decode attended speech envelopes from low-frequency EEG activity. The following sections analyze results with metrics of 1) regression accuracy, 2) classification accuracy, 3) receiver operating characteristic (ROC), and 4) information transfer rate (ITR). Results are shown for each of the regularization schemes, for both forward and backward TRF models. For each regularization scheme, the regularization parameter(s) are tuned to maximize regression accuracy. These parameter values are then used for all regression and classification comparisons. Regression accuracy compares different regularization schemes in predicting test data using the optimal regularization parameter. Classification accuracy uses the regression accuracy values to classify the attended/unattended talker and compares the different regularization schemes in performing this task. The ROC curve visualizes the relationship between the true and false-positive rates for different classifier discrimination function thresholds. Lastly, the ITR describes the impact of decoding segment length on the bit-rate, for different points on the ROC curve.

### 3.1 Regularization Parameter Tuning

The TRF estimation methods, except for the OLS method, use regularization techniques to prevent overfitting and therefore require a selection of the appropriate tuning parameters. Figure 2 shows the correlation coefficient between predicted (validation set) data and the actual target data (*regression accuracy*) over a range of regularization parameters. In general, there is a broad region where validation regression accuracy is flat, which peaks before quickly falling off with increasing *λ*. It is apparent that the regression accuracies obtained with backward TRF models generally are higher than those obtained with forward TRF models.

**Figure 2.**
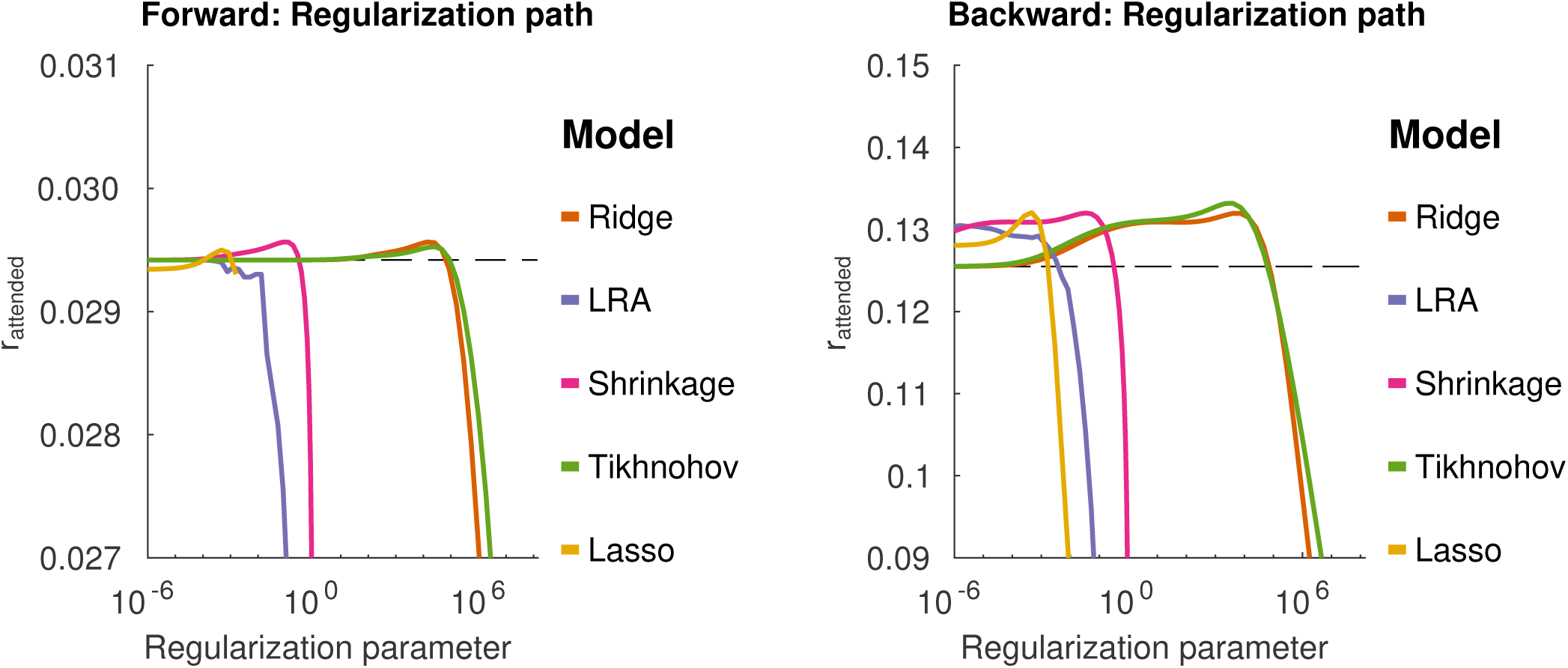
Group-mean validation-set regression accuracies obtained with different TRF estimation methods as the regularization parameters *λ* are varied. The left-hand and right-hand panel present results obtained with forward TRF models and backward TRF models, respectively. The x axis shows the strength of the *λ* regularization parameters. The y axis shows the regression accuracies in terms of Pearson’s correlation coefficients between predicted data and target data. The dashed line shows the regression accuracy for OLS.

Figure 3 shows regression accuracies for TRF models with Elastic Net penalties. Unlike the other linear TRF models investigated in the present study the Elastic Net has two tuning parameters that adjust the balance between *L*1 and *L*2 penalties. This is controlled via the *α* parameter. Similar to the other regularization schemes, for each value of *α*, there is a broad range of *λ* values that give good correlation performance.

**Figure 3.**
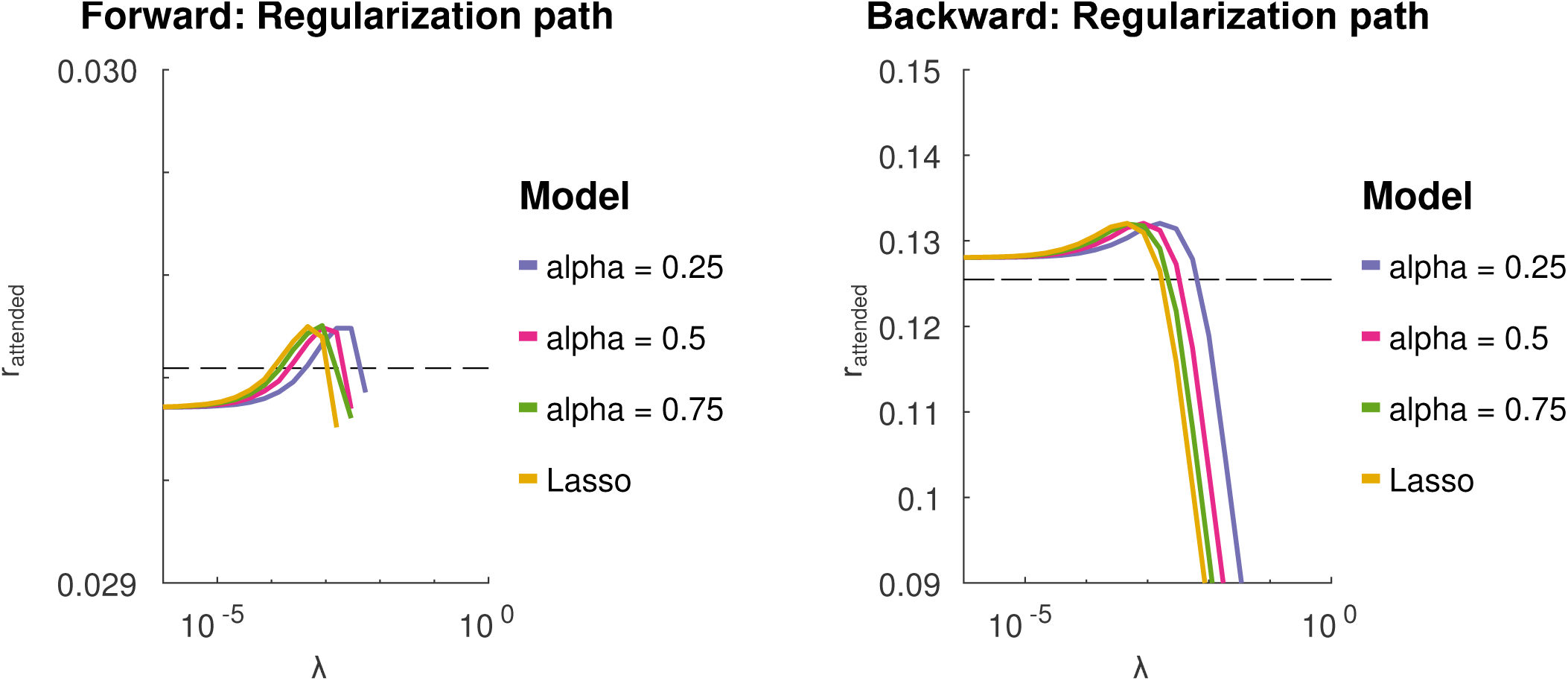
Group-mean validation-set regression accuracies obtained from TRF models with elastic net penalties. The elastic net has to tuning parameters, *λ* and *α*. The two panels show the group-mean validation set regression accuracies cross-validated over a relatively small grid of *λ* and *α* values. The prediction accuracies remain stable over a large range over *λ* values. The dashed line shows the regression accuracy for OLS.

### 3.2 Regression Accuracy

For each regression method (and each value of *α* for elastic net), the TRF model was estimated and the optimum lambda estimated on the training/validation set. This optimal model was then applied to the test set, and the regression accuracy was compared between regression methods. This is shown in figure 4. For forward TRF models, a repeated measures ANOVA with regularization method as the factor (and subject as the random effect variable), found no significant effect of regularization method on the average of correlation coefficients, even when using the average of the correlation coefficients of the 5 channels with the largest correlation coefficients for each subject. For the backward TRF models, a similar repeated measures ANOVA, found a significant effect of regularization method on reconstruction accuracy (*F*_(5,85)_ = 78.0, *p <* 0.01). Tikhonov regularization yielded a regression accuracy that was significantly greater than each of the other schemes, using a Bonferonni correction to account for the family-wise error rate (*p <* 0.05). This is contrary to the expectation that Ridge regression would outperform Tikhonov for the backward model due to the inter-channel leakage introduced by the Tikhonov kernel. Moreover, OLS had a regression accuracy that was significantly smaller than the other schemes (with Bonferonni correction, *p <* 0.01). This highlights the importance of regularization for the backward TRF models.

**Figure 4.**
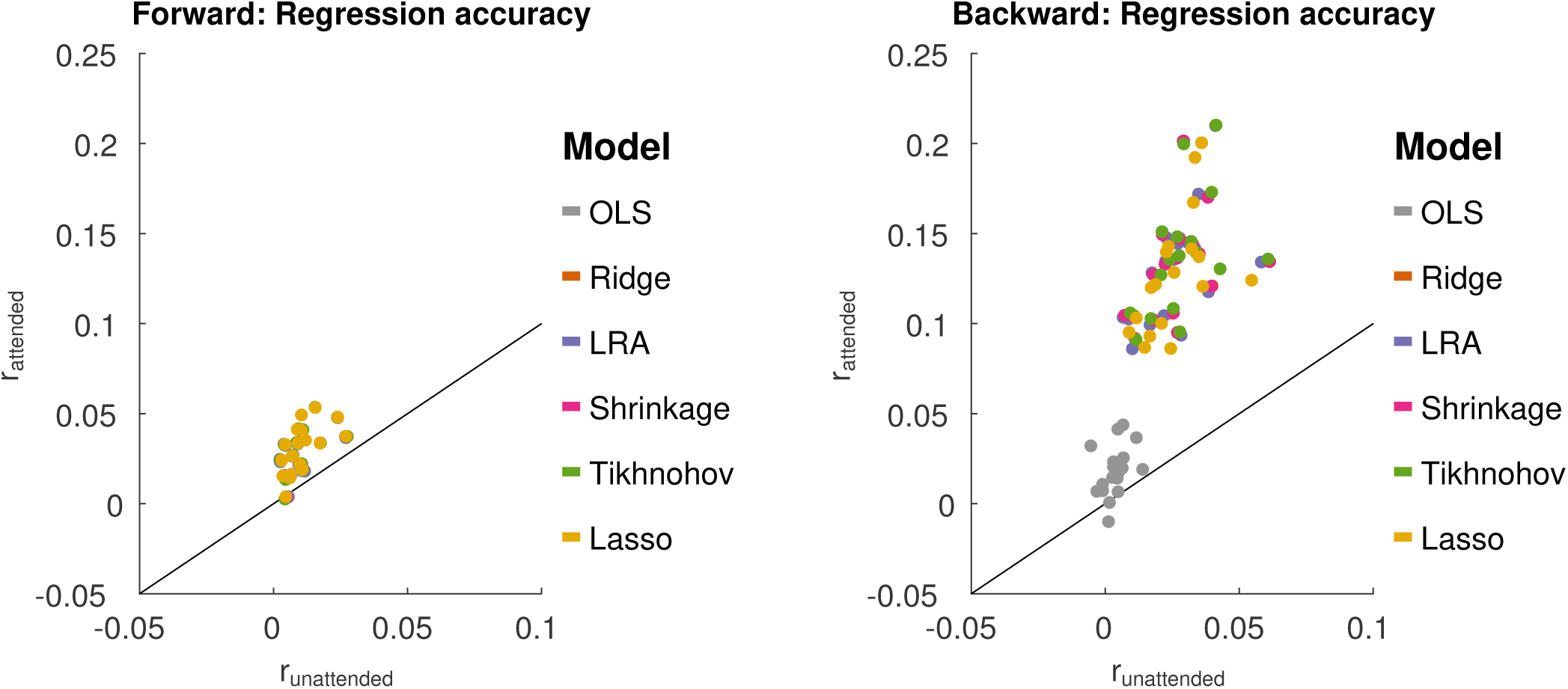
Test set regression accuracies (*r*_*attend*_) for each TRF estimation method plotted against *r*_*unattend*_. Left: results from the forward modeling approach. Right: results from the backward modeling approach. For each scheme (represented by a color), each point represents average data from one subject. The black line shows *r*_*attend*_ = *r*_*unattend*._.

For Elastic Net regularization, *α* values was characterized at 0.25, 0.5, 0.75 and 1 (Lasso) to sample different degrees of sparsity/smoothness. The value *α*=0 (Ridge) was not sampled due to sub-optimal solver performance near this point. A repeated measures ANOVA analysis with factors of *α* and subject, using optimal *λ* values, showed no significant effect of *α* for forward TRF models. This means that adjusting the model sparsity had no significant effect on the reconstruction accuracy. However, a significant effect of *α* was found for backward TRF models (*F*_(3,51)_ = 12.4, *p <* 0.01). A posthoc paired t-test with a Bonferonni correction revealed that the best reconstruction performance was obtained with *α* = 0.25 (*p <* 0.01). It was, however, noted that the average difference between reconstruction accuracies for *α* = 0.25 and *α* = 1 was only 8 *×* 10^−4^.

### 3.3 Classification Accuracy

We further sought to investigate how the different TRF models perform in terms of discriminating between attended and unattended speech on a limited segment of data. The duration of the segment was varied as a parameter (1, 3, 5, 7, 10, 15, 20 and 30s). This was characterized on held-out test data for each TRF method, using the *λ* value that yielded the maximum regression accuracy in the validation data. The results from this analysis are shown in figure 5. A 2-way repeated measures ANOVA with factors of regularization scheme and TRF model (forward or backward), based on 30s decoding segment lengths, found a main significant difference between backward and forward models (*F*_(1,17)_ =17.3, *p <* 0.01), with a significant interaction with the effect of regularization scheme (*F*_(5, 85)_ = 208.9, *p <* 0.01). A posthoc paired t-test showed that backward model performs better than the forward model for all regularization schemes excluding the case where ordinary least squares (OLS) was applied (*T*_17_ = 9.35, *p <* 0.01). For OLS, the forward TRF model outperformed the backward model (*T*_17_ = 7.32, *p <* 0.01).

**Figure 5.**
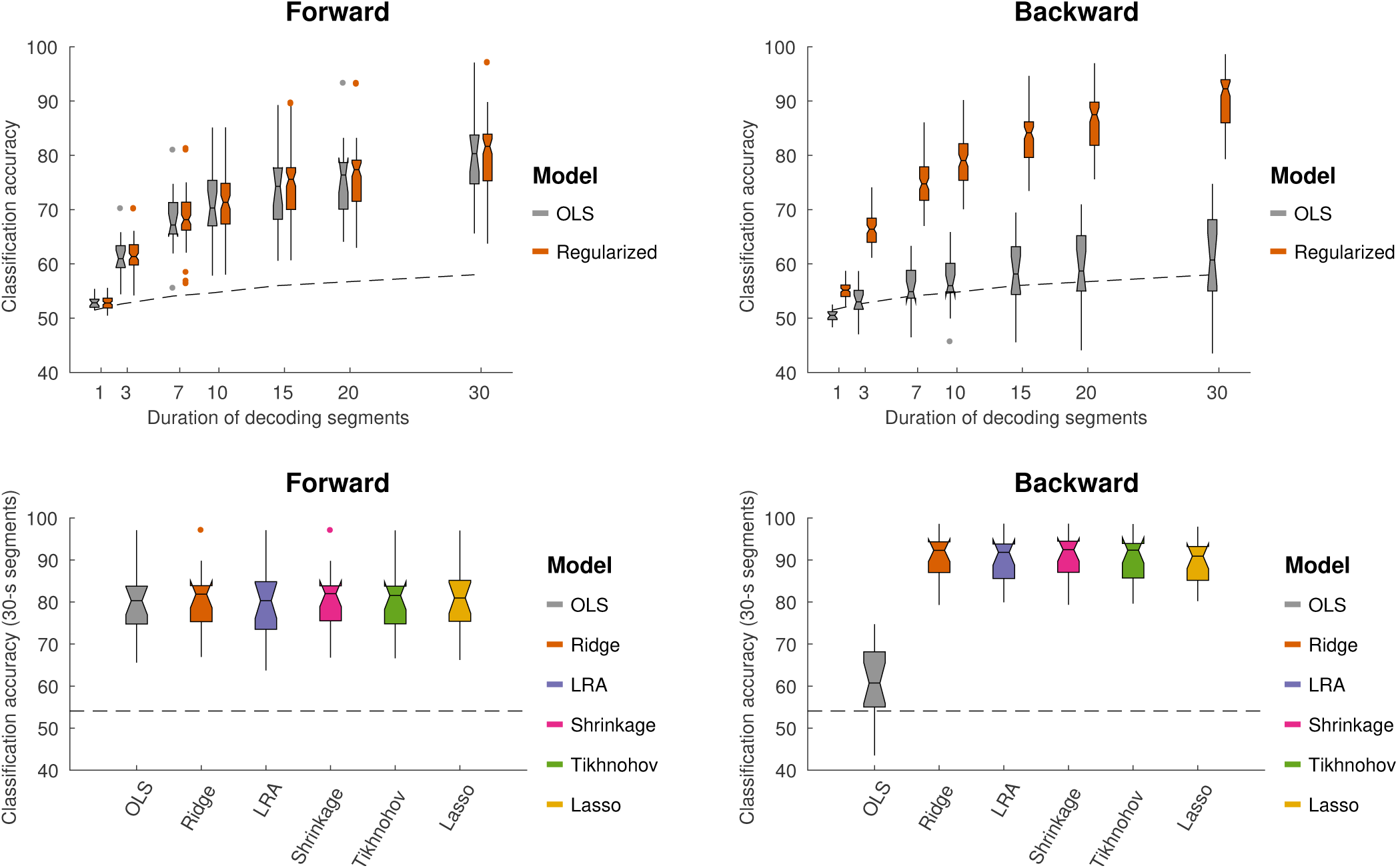
Using different TRF methods to decode selective auditory attention from multi-channel EEG data. Classification performance is shown for different decoding segment lengths (1s, 3s, 7s, 10s, 15s, 20s, 30s). Top-left and -right panels show the classification performance for forward models respectively backward models. Bottom-left and -right panels show the classification performance for 7 s long decoding segments. The different TRF methods are shown in different colors (see legend). Notched boxplots show median, and first and third quartiles. Whiskers show 1.5 *×* IQR. The dashed line shows the above-chance significance threshold at *p* = 0.05.

A repeated measures ANOVA with factors of regularization scheme, applied only to the forward TRF classification accuracy scores, found no significant effect of regularization scheme on classification accuracy. For the backward TRF methods, however, a significant effect of regularization scheme on classification accuracy was found (*F*_(5, 85)_= 229.4, *p <* 0.01). A posthoc paired t-test analysis with a Bonferonni correction revealed that the classification accuracy for the OLS scheme was significantly worse than each of the others *(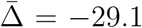*). Lasso performed significantly worse than each of the remaining schemes *(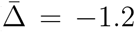*). In short, regularized backward TRF schemes outperform OLS by a relatively large margin, as seen in figure 5.

For Elastic Net regularization, a repeated measures ANOVA with factors of *α* and subject did not find any significant effect of *α* on classification accuracy for forward or backward TRF models.

In summary, for the forward model there was no difference between schemes (regularization and OLS), and for the backward model there was no difference between Ridge, Tikhonov and Shrinkage, but all regression methods were better than OLS.

#### 3.3.1 Relation to regression accuracy

The discrimination between attended and unattended speech streams from EEG data is done in two stages: the computation of regression accuracies, followed by classification. We sought to investigate how the classification accuracies obtained with each TRF model relate to the test set regression accuracies. A plot of this relationship is shown in figure 6.

**Figure 6.**
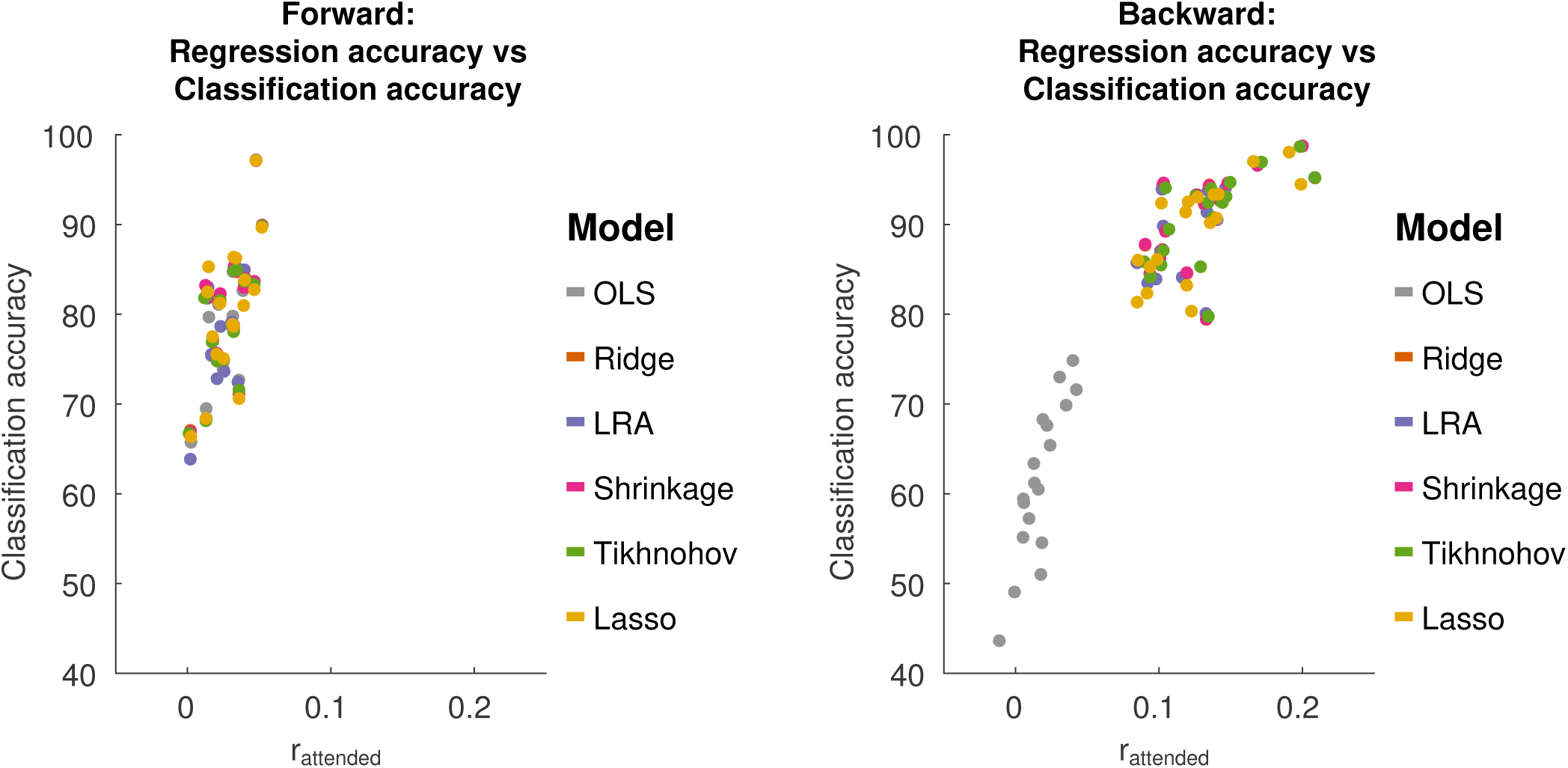
Relationship between regression accuracy and classification accuracy, using 30s decoding segment lengths.

For forward TRF models, the average correlation between regression accuracy and classification performance is 0.69 (*T*_108_ = 9.83, *p <* 0.01), over all regularization schemes. For backward TRF models, the correlation between the regression accuracy and classification performance is 0.89 (*T*_108_ = 22.4, *p <* 0.01). This suggests that classification performance varies with regression accuracy. However, as was previously described for the backward TRF models, while Tikhonov regularization achieved a significantly higher regression accuracy compared to all other methods, it did not achieve a significantly higher classification performance compared to Shrinkage, Ridge Regression or LRA. To explain this, we examined the classification feature in terms of the difference between class means 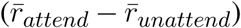 and the within-class standard deviation 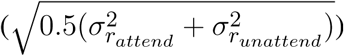. Both of these terms affect the separability between classes.

For backward TRF models, Tikhonov regularization had a significantly larger difference between class means compared to Ridge Regression and Shrinkage (Tikhonov*>*Ridge: *T*_17_ = 1.82, *p* = 0.04), (Tikhonov*>*Shrinkage: *T*_17_ = 1.79, *p* = 0.05). At the same time, the between-class standard deviation was also significantly larger for Tikhonov regularization (Tikhonov*>*Ridge: *T*_17_ = 2.21, *p* = 0.02), (Tikhonov*>*Shrinkage: *T*_17_ = 2.25, *p* = 0.02). This suggests that while Tikhonov regularization yields a better reconstruction accuracy (correlation coefficient), this is offset by an increased variance in the reconstruction accuracy computed over short decoding segments, nullifying any potential gains in classification performance.

### 3.4 Receiver Operating Characteristic

The receiver operating characteristic (ROC) curve, shown in figure 7, shows the relationship between the true-positive rate and false-positive rate for decoding segment trials where the classifier discrimination function lies above a given threshold, as the threshold is varied. The classification accuracy score that we report corresponds to the point on the ROC that lies along the line between (0, 100) and (100, 0). This is also the point at which the Wolpaw information transfer rate (ITR) is estimated, whereas the Nykopp ITR estimation finds a point that lies further left along the ROC curve. The area under the curve is highly correlated with classification accuracy (over all regularization schemes and decoding segment lengths, *r* = 0.99, *T*_862_ =219.9, *p <* 0.01). The Nykopp ITR, on the other hand lies further left along the ROC curve, demonstrating that by avoiding the classification of some trials, it is possible to maximize the ITR.

**Figure 7.**
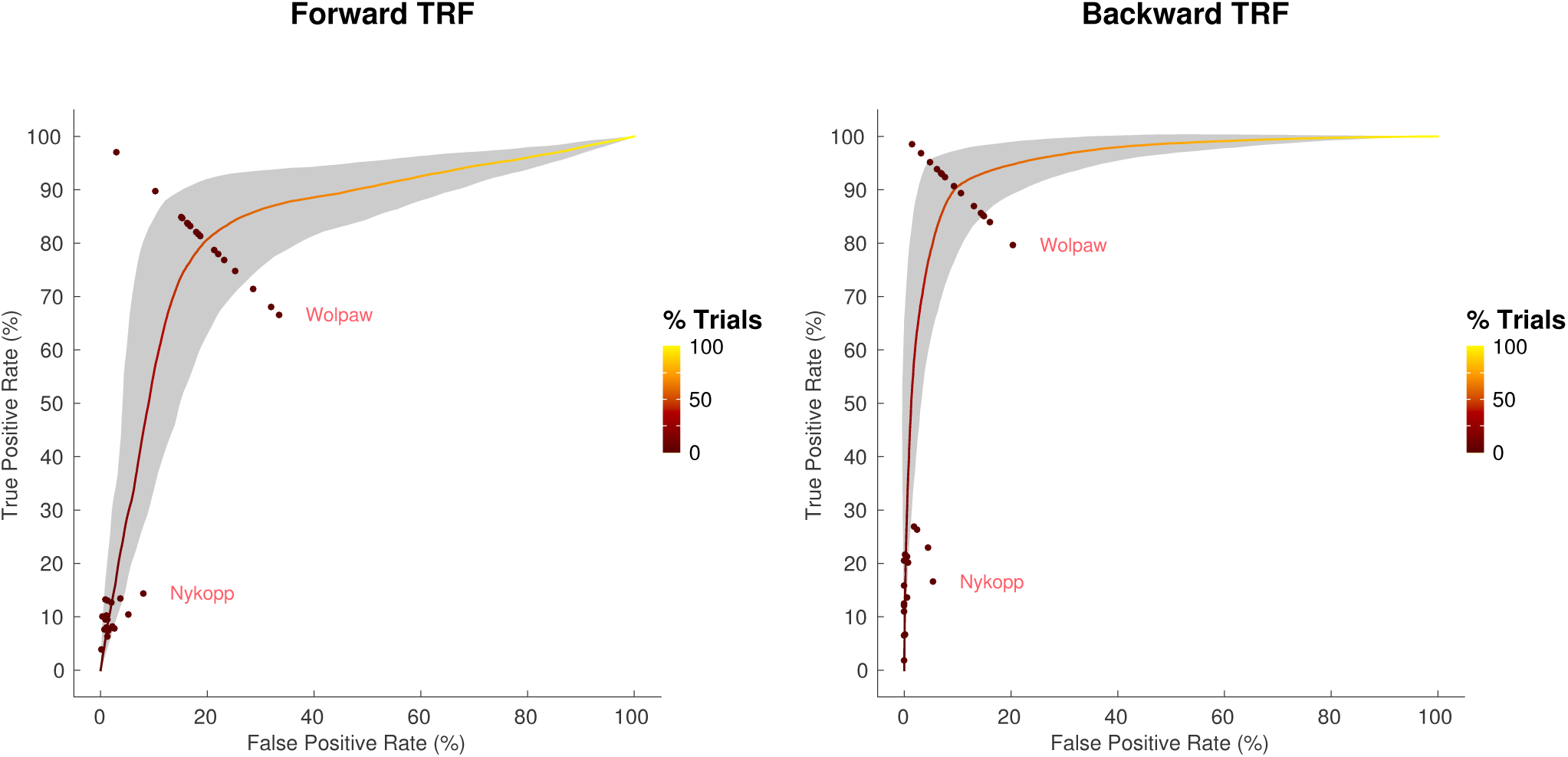
Average receiver operating characteristic curve, with standard deviation band, for 30s decoding segments using Tikhonov regularization. Points at which Wolpaw and Nykopp information transfer rates were evaluated for each subject are shown. Color along curve indicates percentage of decoding segment trials evaluated to obtain each point. The gray band indicates the standard deviation boundaries of the curve in both x and y directions.

### 3.5 Information Transfer Rate

The Wolpaw ITR represents the transfer rate when all decoding segments are classified, whereas the Nykopp ITR represents the maximum achievable transfer rate when some classifications are withheld based on classification discrimination function output. Figure 8 shows the Wolpaw and Nykopp ITR values as a function of decoding segment duration, based on TRFs computed with Tikhonov regularization. Both the Wolpaw and Nykopp ITR show an increase followed by a decrease with increasing decoding segment duration. The plots suggest that for brain computer interface applications with fixed decoding segment lengths, it may be advisable to use decoding segments of 3-5 seconds to maximize the ITR. While the Nykopp measure is an upper-bound, its increase over the Wolpaw ITR value (forward TRF, 5s: *T*_17_ = 13.1, *p <* 0.01), (backward TRF, 5s: *T*_17_ = 16.7, *p <* 0.01) demonstrates that by adjusting the classifier decision function cutoff, it could be possible to increase the ITR.

**Figure 8.**
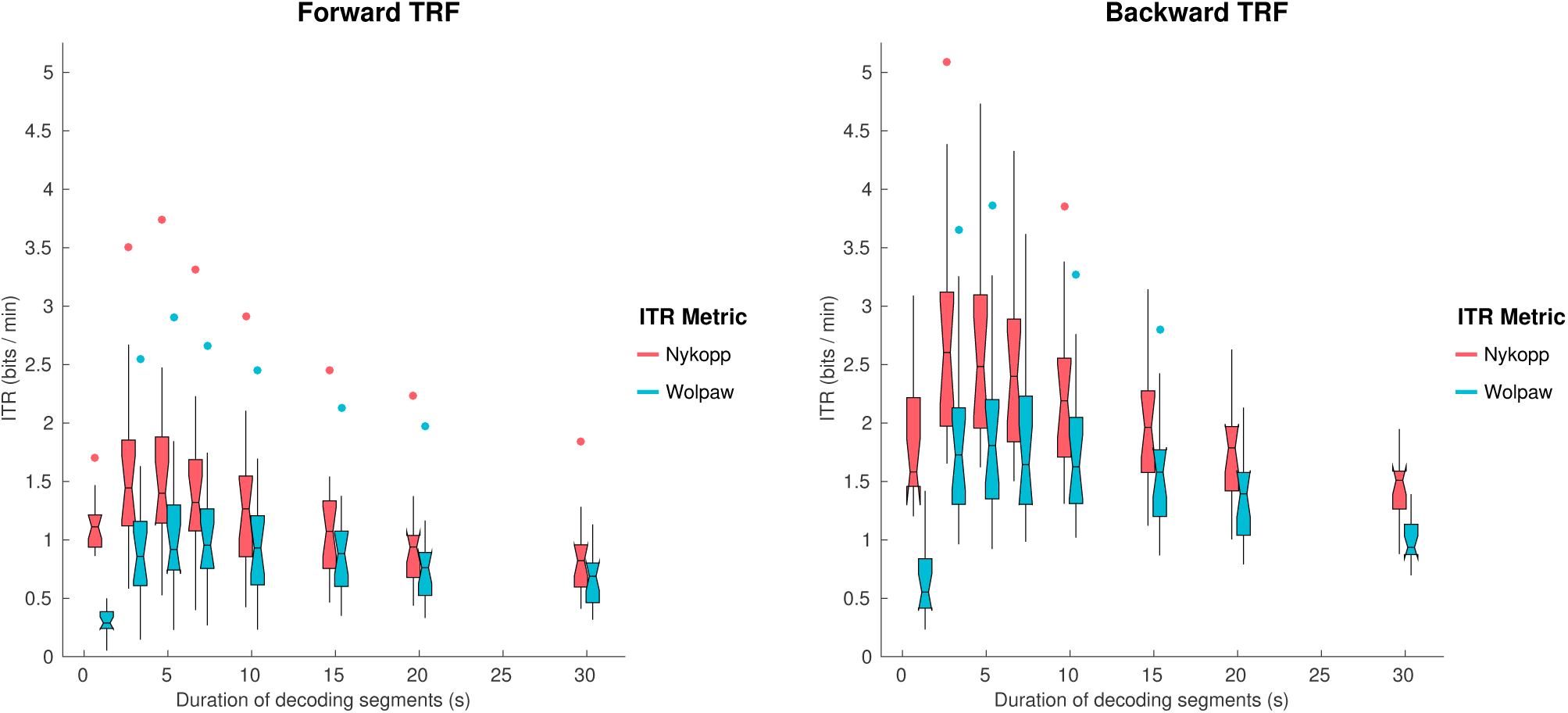
Wolpaw and Nykopp information transfer rates (ITR) as a function of decoding segment duration for the forward and backward TRF models, using Tikhonov regularization.

## 4 DISCUSSION

In this study, we systematically investigated the effects of TRF estimation methods on the ability to decode and classify attended speech envelopes from single-trial EEG responses to speech mixtures. The performance of stimulus/EEG decoders based on forward TRF models (mapping from attended speech envelopes to multi-channel EEG responses) and backward TRF models (mapping from EEG response back to speech envelopes) were compared. It was found that the backward TRF models outperformed the forward TRF models in terms of classification accuracies. We hypothesize that TRF models do a better job of predicting audio (the backward model) than EEG data (the forward model) because the EEG data contains a lot of information from other brain functions. It is simpler to filter out these signals, as is done in the backward model, than it is to predict them (as the forward model would need to do to achieve higher correlation). Different regularization schemes were not found to significantly affect the forward TRF classification accuracies. However, for the backward TRF models, the decoding schemes that yielded the best classification accuracy were Ridge Regression, LRA, Shrinkage and Tikhonov. Lasso had a lower classification accuracy by a small but significant margin. Classification accuracy increased monotonically as a function of duration, reflecting the greater amount of discriminative information available in longer segments. ITR however peaked at an intermediate segment duration, reflecting the tradeoff between the accuracy of individual classification judgments (greater at long durations) and number of judgments (greater at short durations). The optimum was around 3-5s.

For the analysis, we used different linear approaches to decode selective auditory attention from EEG data. These analyses all relied on the explicit assumption that the human cortical activity selectively tracks attended and unattended speech envelopes. To fit the models, we made a number of choices based on common practices in literature, and with the goal of being able to compare TRF methods. For example, a 500 ms TRF kernel used as was done by others (14). While shorter kernels have been explored as well (23), a longer one tests the ability of the TRF method to handle a larger dimensionality and allows for a more flexible stimulus-response modeling capturing both early and late attentional modulations of the neural response. Additionally, we chose to focus on 1-9 Hz EEG activity as the attentional modulation of EEG data has been found prominent in this range. It is likely that other neural frequency bands robustly track attended speech (e.g. high gamma power (25)) and that the neural decoders potentially could benefit from having access to other neural frequency bands. This is, however, outside the scope of this paper.

### 4.1 Decoding selective auditory attention with forward and backward TRF models

The forward TRF models performed significantly worse than the backward TRF models in terms of classification accuracies. Single-trial scalp EEG signals are inherently noisy, in part because activity picked up by each electrode reflects a superposition of activity from signals that are not related to the selective speech processing (3). We refer here to any aspects of the EEG signals that systematically synchronize with the attended speech streams as target signals and anything that does not as noise. To improve the signal-to noise ratio one can efficiently use spatio-temporal filtering techniques. This in part relates to the fact that stimulus-irrelevant neural activity tends to be spatially correlated across electrodes. The spatio-temporal backward models implicitly exploit these redundancies to effectively filter out noise and improve signal-to-noise-ratio. This makes them fairly robust to spatially correlated artifact activity (e.g. electro-ocular and muscle artifacts) when trained on data from a large number of electrodes. This is also reflected in the high classification accuracies that were obtained with the backward models. However, for the relatively high number of electrodes used in this study, it was found that the spatio-temporal reconstruction filters were effective only when properly regularized.

The forward models, on the other hand, attempt to predict the neural responses of each electrode in a mass-univariate approach. These models do not, therefore, explicitly use cross-channel information to regress out stimulus-irrelevant activity. The relative contribution of the individual channels to the classification accuracies were instead found via an SVM trained on correlation coefficients computed per channel, over short time segments. It can therefore be beneficial to apply dimensionality reduction techniques (e.g. independent component analysis (2) or joint decorrelation (6)) to represent the EEG data as a linear combination of fewer latent components prior to fitting the forward models. Alternatively, canonical component analysis can be used to jointly derive spatio-temporal filters for both audio and EEG such that the correlation between the filtered data is maximized (7).

#### 4.1.1 Regularization

Each regularization scheme makes certain assumptions and simplifications that are therefore adopted by studies employing them. Because these methods have not been previously evaluated side by side, it is unknown how valid these assumptions are.

While no regularization (OLS) was found to work well for forward TRF models in producing classification accuracies roughly in line with regularized models, this method performs relatively poorly when applied to backward TRF models. This is likely reflective of the higher dimensional TRF kernel required for the backward problem. For comparison, a forward TRF model had 33 parameters (per channel) that needed to be fit, whereas a backward TRF model had 2, 178 parameters.

We generally found that the reconstruction accuracies (*r*_*attend*_) plateaued over a large range of *λ* values for linear TRF models (Figure 2). In fact, fixing the regularization parameter to a high value did not strongly affect the decoding accuracies compared to doing a hyperparameter search (this was tested with ridge regression with a fixed large *λ* value).

Elastic net regularization permits the adjustment of the balance between L1 and L2 regularization via the *α* parameter. For the backward TRF model, it was shown that a smaller *α* value improved the correlation between the reconstructed and attended audio stream by only a narrow margin. The *α* value had no significant impact on classification accuracy for either forward or backward TRF models. As such, the higher classification performance of Ridge Regression (*α* = 0), compared to Lasso (*α* = 1) may be a result of differences between solvers (MATLAB’s *mldivide* versus GLMNET (30)).

For the forward model, all regularization schemes yielded reconstruction and classification accuracies that were not significantly different from each other. For the backward model, Tikhonov regularization yielded the best regression accuracy. However, it was found that this did not lead to a better classification accuracy compared to other L2-based regression schemes (i.e. Ridge, Shrinkage and LRA) due to an associated increased variance in the correlation coefficient computed over short decoding segment lengths. It has been reported that, in practice, the Ridge Regression approach appears to perform better than LRA (33). While LRA yielded marginally lower mean regression accuracy and classification performance than Ridge Regression, this was not found to be significant. LRA removes lower variance components after the eigendecomposition of **X**^*T*^ **X**, essentially performing a hard-threshold. In contrast, Ridge Regression is a smooth down-weighting of lower-variance components (3).

### 4.2 Realtime Performance

The information transfer rate results provide insight into how classification performance can be optimized. It is worth noting that the ITR measures represent particular points along the ROC curve, as is illustrated in Figure 7. For a binary classification problem, with balanced classes, the Wolpaw ITR corresponds to the point on the ROC curve along the line connecting the corners of the plot at coordinates (100, 0) and (0, 100). The Nykopp ITR, on the other hand corresponds to the point that maximizes the ITR, essentially trading the number of classified samples for increased classification accuracy. In practice, other considerations besides ITR can influence the choice of the point on the ROC. For instance, if there is a high penalty on incorrect classifications, then the classifier threshold may be adjusted to operate at another point on the ROC curve. In short, the ROC and ITR are useful tools in identifying a suitable balance between sensitivity and specificity.

The ITR results in the present study suggest a 3-5 s decoding segment length to achieve the maximum bit-rate. It should be noted that this assumes that switches in attention can occur frequently, on the order of the decoding segment length. In cases, where switches in attention are known to be sparse *a priori*, it may instead be more desirable to increase decoding segment length and sacrifice bit-rate to put more emphasis on accuracy, since the loss in bit rate due to long decoding segments is only evident during attention switches. Such an approach was taken by O’Sullivan and colleagues (24), where the theoretical performance of a realtime TRF decoding system was characterized for switches in attention every 60 s. In that study, a decoding segment length between 15-20 s was reported as optimal to achieve the best speed-accuracy tradeoff.

### 4.3 Summary

There are many methods that can be used to compute TRFs. The present study uses a baseline dataset and procedures for the evaluation of these TRF methods. In consideration of the multiple applications in which TRF functions are used, primarily dealing with reconstruction accuracies or classification performance, this paper considered multiple metrics of TRF performance. By characterizing the regularization and performance of the TRF methods, and the relationship between performance metrics, a more complete understanding of the validity of the assumptions underlying each TRF method is provided, as well as the impact of the assumptions on the end result. While these experiments were done with EEG data, we expect that the results apply equally to magnetoencephalography (MEG) data. The key findings from this study were 1) the importance of regularization for the backward TRF model, 2) the superior performance of Tikhonov regularization in achieving higher regression accuracy although this does not necessarily entail superior classification performance, and 3) optimal ITR can be achieved in the 3-5 s range and by adjusting the classifier discrimination function threshold.

## 5 ACKNOWLEDGEMENTS

This work was supported by the EU H2020-ICT grant 644732 (COCOHA), and grants ANR-10-LABX0087 IEC and ANR-10-IDEX-0001-02 PSL. It draws on work performed at the 2015 and 2016 Telluride Neuromorphic Engineering workshops.

## Footnotes

1 http://www.ine-web.org/software/decoding

2 http://www.ine-web.org/software/decoding

3 http://www.cocoha.org/the-cocoha-matlab-toolbox

